# Sub-daily Bermuda Atlantic Time Series virus sampling reveals taxonomy, host, and functional differences at the population, but not community level

**DOI:** 10.1101/2025.10.20.683370

**Authors:** Alfonso Carrillo, Emily Hageman, Lauren Chittick, Anna I. Mackey, Kimberley S. Ndlovu, Funing Tian, Naomi E. Gilbert, Daniel Muratore, Dean Vik, Gary R. LeCleir, Christine Sun, Ho B. Jang, Ricardo R. Pavan, Joshua S. Weitz, Steven W. Wilhelm, Matthew B. Sullivan

## Abstract

Ocean microbes contribute to biogeochemical cycles and ecosystem function, but they do so under top-down pressure imposed by viruses. While viruses are increasingly understood spatially and beginning to be incorporated into predictive modeling, high-frequency ocean virus dynamics remain understudied due to methodological challenges. Here we sampled stratified Bermuda Atlantic Time Series (BATS) waters for 112 hours at sub-daily 4-(surface) or 12-(deep chlorophyll maximum) hour intervals, purified viral particles from these samples, sequenced their metagenomes, and used the resulting data to characterize high-frequency virus community dynamics. Aggregated community diversity metrics changed with depth, but were not statistically significant temporally at a fixed location. However, finer-scale population-level analyses revealed both depth and temporal change, including physicochemical depth-driven differences and, in surface waters, thousands of viral populations that exhibited statistically significant diel rhythms. Statistical analyses revealed three main archetypes of temporal dynamics that themselves differed in abundance patterns, host predictions, viral taxonomy, and gene functions. Among these, highlights include viruses resembling an archetype with a night peaking pattern in activity that include an over-representation of viruses that putatively infect *Prochlorococcus*, a phototrophic cyanobacteria. Together, these efforts provide baseline community-and population-scale short-time-frame observations relevant to future climate state modeling.

## Introduction

The oceans play key roles in global biogeochemical cycles, including buffering against human-accelerated climate change [1]. However, this buffering is self-limiting - ocean biogeochemistry is directly impacted by climate change as warming surface waters increase seasonal stratification [2] and see reduced nutrient concentrations. These impacts are particularly important over the vast oligotrophic ocean regions that cover approximately 30-50% of the Earth’s ocean surface [3,4]. Such altered surface ocean functions are hypothesized to impact the ocean’s ability to absorb atmospheric carbon dioxide and sink surface-produced carbon to the deep sea via the biological carbon pump. While one-third of anthropogenic carbon dioxide released into the atmosphere is absorbed into surface waters via mass action [1], its fate is dictated by plankton – including bacteria, archaea and microbial eukaryotes that serve as the base of the food web and drive the biological carbon pump.

Viral roles in these ocean biological carbon pump processes are recently being revisited. For decades, the viruses that infect these cells were thought to keep carbon in the dissolved phase by lysis resulting in remineralization – a process known as the viral shunt [5]. However, viral lysis may also generate sticky aggregates that sink out of the photic zone. This ‘viral shuttle’ has been hypothesized as a mechanism that reduces the retentiveness of the microbial loop [6–10]. Recent work leveraging machine learning, statistical modeling, and global ocean datasets from the *Tara* Oceans expeditions provides evidence in support of this hypothesis given that globally ocean carbon flux is best predicted by viral abundances – even more so than abundances of bacteria, archaea, or eukaryotes [11]. In parallel, viral infection of microbial hosts has shown viruses to reprogram infected cells (termed ‘virocells’, [12]) into entities that are metabolically and biogeochemically different from their uninfected sister cells [13–16]. This reprogramming can also apply when cells mutate to successfully defend themselves against virus attack. Towards this, recent experimental work shows spontaneous virus-resistant marine bacterial mutants alter their carbon substrate utilization, metabolite secretion, and aggregation and sinking rates in ways that will drastically alter these cells’ ecosystem inputs and outputs [17]. Thus, viruses can alter a cell’s ecosystem outputs well beyond simple lysis.

Early work on Kill-the-Winner models showed how viruses infecting a single microbial host could enable fluctuating host diversity through negative frequency dependent selection [18–20]. These models have been extended to include increased complexity - spanning interaction networks, community dynamics, and feedbacks with ecosystem functioning [21–29]. However, a major obstacle to extending these models to the Earth System scale is the lack of observational data to guide model incorporation, with sampling challenges resulting in relatively little being known about marine microbial and viral temporal dynamics. Some time series sampling efforts have assessed, at monthly resolution, abundance changes in bacteria and archaea (using PCR-amplicons and/or metagenomic sequencing), and picoeukaryotes (via flow cytometry) – and this has been done for years (e.g., 7 year Bay of Banyuls [30]) or even decades (e.g., >30 years Hawaii Ocean Time (HOT) Series [31]). However, these studies largely excluded viruses, except as bycatch in early low-resolution metagenomic surveys [32,33]. Other time series similarly surveyed bacteria, archaea, and picoeukaryotes over long time scales monthly, while also including viruses (e.g., >15 years at San Pedro Ocean Time (SPOT) Series [34–36]).

Although they are based on amplicon-based approaches [37], these studies nonetheless provide strong evidence of seasonality whereby bacteria, archaea, and picoeukaryote abundances cycle yearly and then return to (nearly) the same state with only slight year-to-year baseline shifts.

At a much finer temporal resolution, some studies have explored diurnal rhythms in ocean systems for microbes and their viruses. Sub-daily timescale metatranscriptome-derived microbial and viral transcript measurements have helped identify strong diel rhythms of gene expression along with temporal relationships between gene expression and metabolite concentration [38–42]. Observations of transcript activity between viruses and their hosts have shown that viruses can often coordinate their activity with host diel cycles, responding to environmental cues in synchrony with their hosts [40,41]. Other work has taken more targeted approaches by focusing on one group, cyanobacterial viruses (or cyanophages), and revealed transcriptional rhythms and adsorption rates in the field that laboratory studies then linked to diurnal photosynthetic activity [43]. Along with this, amplicon-based marker gene fragment patterns (e.g., terminal restriction fragment length polymorphism) have similarly been used to target T4-like viruses to reveal daily dynamics over the course of 38 consecutive days that correlate virus operational taxonomic units (vOTUs) with bacterial OTUs to document short-term variation in both microbial and viral communities [35]. Together these studies demonstrate clear sub-daily dynamics, but are limited in that they again either catch viruses as a byproduct of microbial sampling (e.g., prokaryotic metatranscriptomics) or target specific virus groups (e.g. cyanophages or T4-like viruses) rather than community-wide signals often doing so with highly degenerate primer strategies that might confound quantitative data generation.

In the decade since these studies, viral metagenomics [37,44] and our understanding of virus population biology [45–47] have matured to the point that there is opportunity to study viruses at increasingly high taxonomic and temporal resolution, even over a short timescale. The Bermuda

Atlantic Time Series (BATS) – a sentinel model ecosystem for “future oceans” due to its increased stratification and acidification [48] – has been surveyed via DNA staining and epifluorescence microscopy to quantify virus and microbe abundances and found seasonality over a decade of sampling from 2000-2009 [49]. While BATS samples were targeted by these extensive SYBR-stained virus abundance data and early sequencing technology was applied to a single virus sample [50], no genome-resolved virus sequencing data were available at BATS. This prevents ecological inferences like the kinds of viruses, their hosts, their functions, and their metabolic reprogramming capacities. Towards this, recent work employed time-resolved metagenomics at BATS and then compared viral population-based abundances inferred from those captured in paired cellular and viral fraction metagenomes. This revealed that viral fraction population data represents the integral of cellular fraction infections and multiple days of viral turnover [51], but left open questions about viral community and population-level changes through time.

Here we build upon these prior efforts by establishing a 112-hour time series - sampled every 4 hours in surface waters and, every 12 hours at the deep chlorophyll maximum - of virus metagenomic data at BATS during late seasonal stratification as a proxy for future climate-change-impacted oceans. This resulted in detailed, sub-daily insight into community-and population-level viral dynamics for 48,428 viral populations at BATS.

## Results

### Assembly of a high quality and detailed viral reference database at BATS

To estimate sub-daily virus population-level dynamics, we established a high-resolution dataset from surface (SUR) and deep-chlorophyll maximum (DCM) waters sampled at BATS between October 12-17, 2019 (**Fig 1A-C**). A total of 39 virus concentrates were prepared via chemical flocculation [52] (see Methods) of samples collected every 4 (SUR waters) or 12 (DCM waters) hours over a 112-hour time course (**Fig 1C**). To follow the same parcel of water through the time-course, sampling was done in a Lagrangian manner using a surface buoy with an underwater drogue (at 30m depth) to ‘track’ the water for the length of the study. Sampling started at 16:00 local time (GMT-3) and depths were determined with each collection from water column profiles of temperature and salinity collected by the ship’s CTD system as described previously [53].

**Figure 1.**
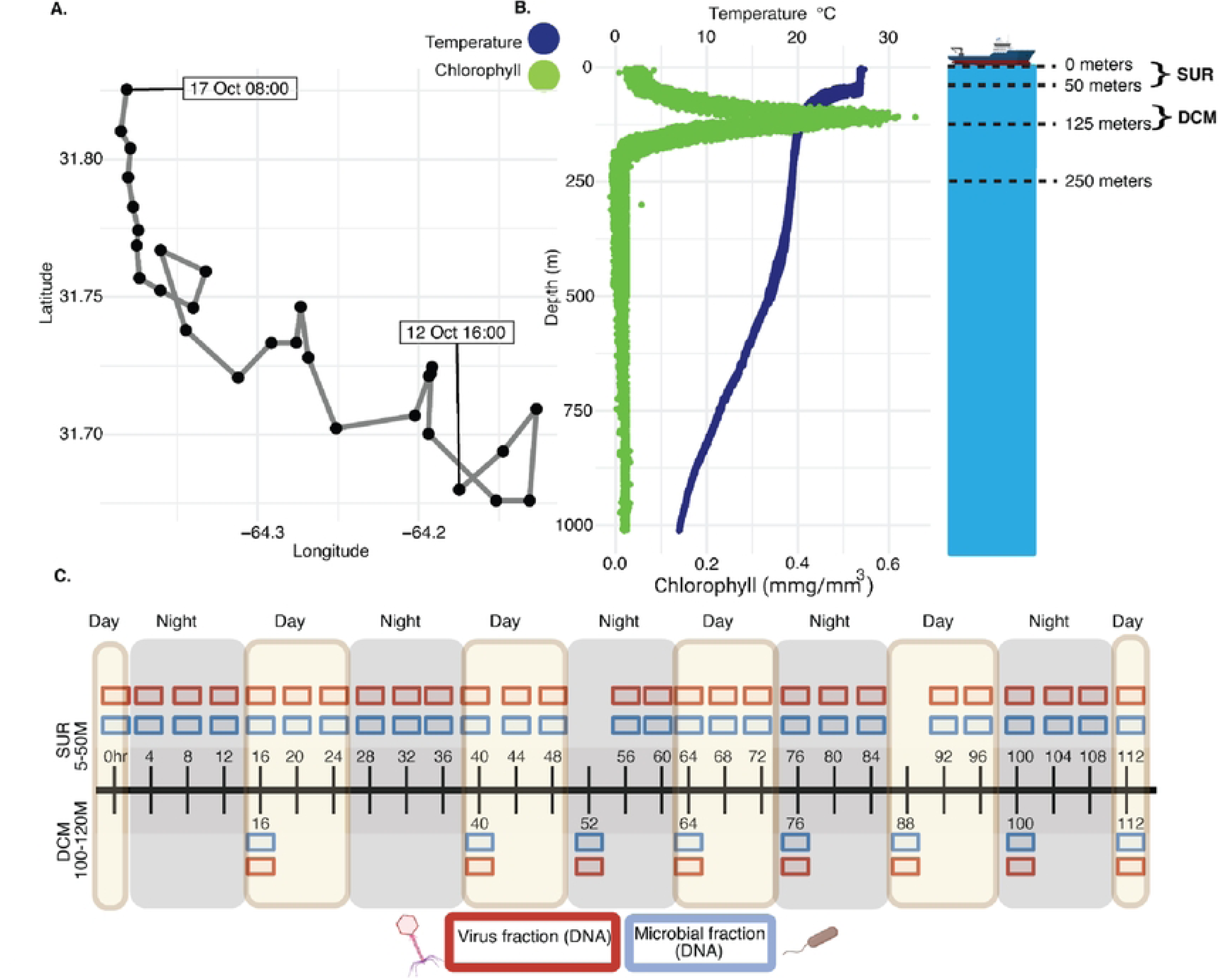
Field site location, depth profiles, and sampling schematic. **A.** The Bennuda Atlantic Time Series is located in the Sargasso Sea portion of the Atlantic Ocean. Lagrangian sarnpling track for data collection is shown with each black point representing a CTD-cast starting on Oct. 12^th^, 2019 at 16:00 local tin1e (GMT-3) and ending on Oct. 17’1\ 2019, at 08:00. **B.** Depth profile of chlorophyll (green) and temperature (blue) down to 1000 meters depth, and ship sche1natic to emphasize key depths in the water colunu1 including ship (On1), surface (0-501n), and DCM (105-120m). **C.** Sampling schernatic to show the depth (surface or SUR versus deep chlorophyll rnaximum or DCM), frequency (every 4 and 12 hours for surface and DCM, respectively), size fraction (virus red boxes or cellular blue boxes), and tirne of day (yellow or gray background shading) of each sample taken.

The resultant virus-fraction concentrates were resuspended in a buffer, DNA extracted, sequencing libraries that targeted double-stranded DNA viruses only were prepared, and then short-read sequenced (see Methods) to an average depth of 147M reads per sample. The resultant 39 virus metagenomes were assembled (see Methods) into 1.48B contigs (≥10kb) and 228,013 of these were conservatively identified as viral (using the VirSorter 2 Standard Operating Procedure VS2_SOP [54,55]). These virus contigs were then clustered into 48,428 viral populations using community consensus cut-offs of 95% average nucleotide identity over 80% of the contig coverage [46,47,56] with an average genomic length captured of 10.28kbp. While other ocean datasets reflect more dispersed sampling schemes, such as the deeply sequenced GOV2 dataset that spread its sequencing across 79 sites [46], here we provide a more focused sampling of the BATS site to maximally identify virus populations at this site. Indeed, this provides us with many more captured viruses at a single site (compare 48,428 with an average of 2,083 viral populations per site in GOV2), at a similar genomic coverage per population (compared 10.28kb versus GOV2’s average genomic length of 10.17kbp [46]).

Taxonomically, these populations are thought to represent approximately species-level taxa as prior studies inferred from (i) gene flow, selection, and genomic fixation indices [45], (ii) host range based phenotypic variation in heterotrophic viruses [57], and (iii) natural breaks in sequence similarity for nearly 500K virus populations in GOV2 [46].

### Community level diversity significantly varies with depth, but not through 112-hour time-course

We first sought to assess how community-level diversity changed with depth and across our high-frequency time course (see Methods). For depth-resolved comparisons, we hypothesized that viral diversity would differ significantly between surface (SUR - sampled from 5m) and deep chlorophyll maximum (DCM) samples, due to the inherent differences in physicochemical features such as temperature, and light penetration that also impact biology (e.g., chlorophyll as a proxy for phototrophic biomass) [[58–60]; **Fig 1B, Fig S1**]. Indeed, we found that virus community alpha diversity (Inverse Simpson’s Index) was significantly higher in SUR versus DCM waters (Wilcoxon Rank Sum *P* = 0.000028) (**Fig 2A**), and this remained stable for nearly all time points throughout the 112-hour time course (**Fig 2A**). These findings complement other daily observations at BATS where extracellular viral abundances had clear depth-specific differences inside and outside the surface mixed layer [51]. Additionally, we evaluated Inverse Simpson’s diversity changes within SUR versus DCM depths of water columns sampled throughout the global oceans alongside our own. This revealed that BATS had higher alpha diversity in SUR as opposed to the DCM, contrary to other previously sampled regions ([46], **Fig S2**).

**Figure 2.**
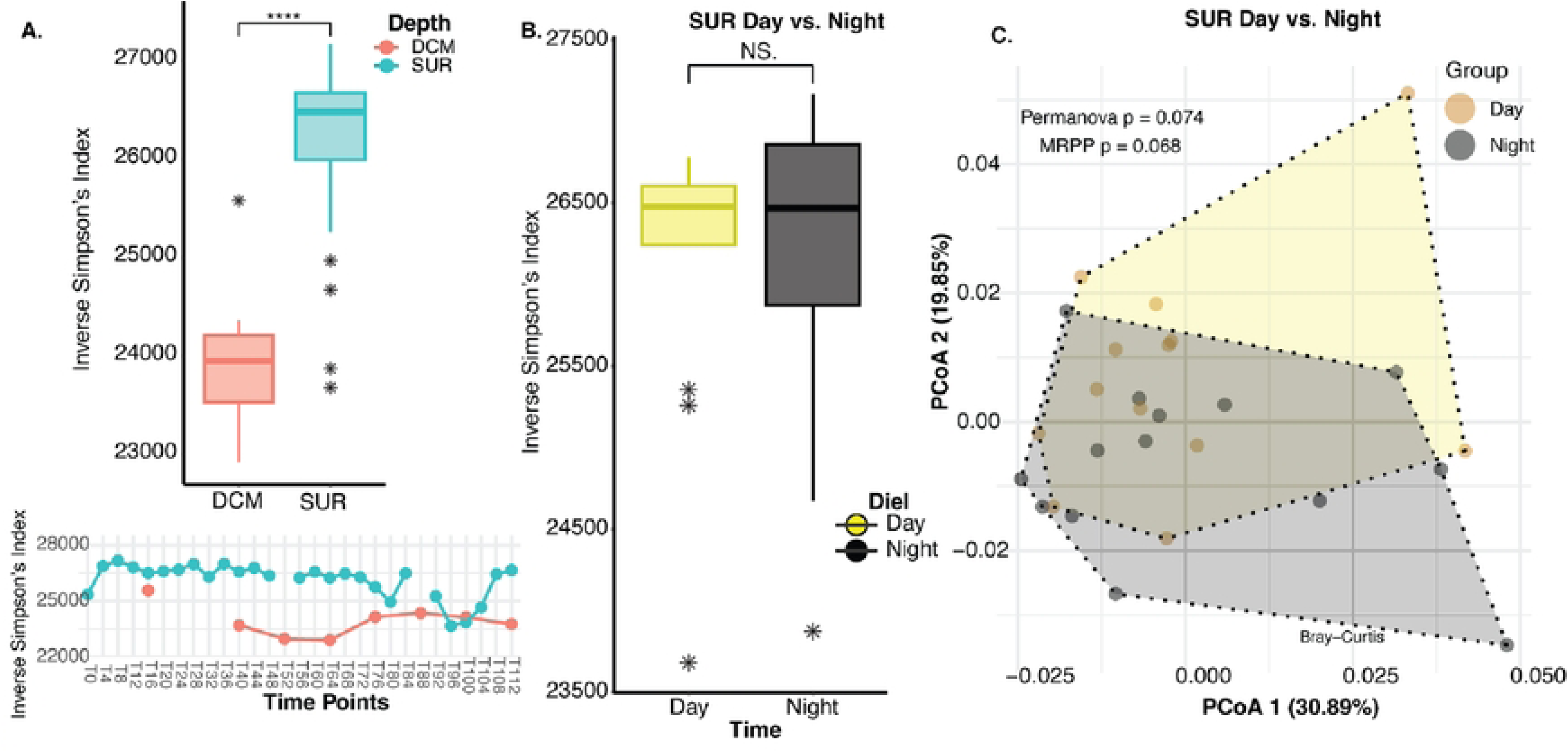
Corn1nunity-level statistics across,vater depths and tin1es of sampling. **A.** Inverse Sitnpson’s Index box plot showing that vi1us co1rununity diversities were statist.ically significantly different (Wilcoxon Rank Sum *P* = 0.000028) between SUR (blue points) vs. DCM (pink points). Line graph below depicting alpha diversity (Inverse Si1npson’s) changes across time points for SUR (blue poit1ts) and DCM (pitik points) samples. **B.** Inverse Simpson’s Index box plot of day (yellow points) vs. night (black points) for SUR san1ples showing that vitus conununity diversities were not significantly different (Wilcoxon Rank Su1n *P* = 0.62, denoted as NS m the figure). C. Principal coordinate analysis of Bray Curtis dissimilarity distances also shows that day versus night virus co1n1nunity dissimilarities were not significantly different (Multi-Response Permutation Procedure, or MRPP, p=0.068). Shaded polygons outline day and night groups.

We also assessed the impact of the diel cycle on diversity proxies in the surface samples only.

Specifically, we evaluated how alpha-and beta-diversity metrics varied between nighttime (20:00) and daytime (08:00). Although some microorganisms possess unique circadian rhythms and dependencies on the light-dark cycle [61], community-wide virus diversity metrics were not significantly different whether inferred at the level of alpha diversity (Inverse Simpson’s diversity metric; Wilcoxon Rank Sum *P* = 0.62; **Fig 2B**) or beta diversity (PCoA ordination of Bray-Curtis dissimilarity measures; MRPP *P* = 0.068; **Fig 2C**). Thus even though day/night light intensity and microbial taxa change during this 112 hour sampling [53], our findings show that aggregate viral community diversity measures do not change. Our 2019 BATS viral sampling expands upon prior 2017 BATS viromic sampling [51] by increasing resolution (4-vs 12-hour sub-daily time scale) and adding 20x fold more viruses at a single site (48K vs 2.3K). We reasoned then that these significant dataset improvements provided a better opportunity to assess high-resolution virus community dynamics at BATS.

### Viral populations and potential metabolic gene repertoire vary across depth

Given that the viral depth-related differences shown above were explored at the community-level, we continued by exploring depth-related changes at the virus population level. First, we predicted hosts for our 48,428 virus populations to assess their patterns across the two depths sampled. Notably, we used aggregate *in silico* prediction strategies (see Methods) and maximized our ability to predict hosts (prior benchmarking suggests host prediction improvement of 25% for most systems [62] by augmenting the standard database available with 89 high-quality metagenome-assembled genomes (MAGs) from prior BATS work [51]. This resulted in predicted hosts for 11,814 (24.39%) of our 48,428 total populations, a large improvement on the 6 (0.26%) of 2,301 total viral populations whose hosts could be predicted in that prior study [51].

SUR versus DCM depth comparisons revealed some populations were shared across depths, whereas others exclusive. As expected, hosts whose virus populations were better sampled displayed abundance changes across the depths. For example, viruses that infect heterotrophic bacteria such as *Pelagibacter* (SAR11 clade I), *Pelagibacter_A* (SAR11 clade II), and *AG-337-I02* (SAR11 clade V) were among the most represented host predictions in SUR, with *AG-337-I02* and *Pelagibacter* ∼4-fold or higher in abundance at the surface compared to DCM (**Fig 3A-B)**. This SUR enrichment signal of SAR11 clade infecting viruses parallels SAR11 bacteria being one of the most abundant heterotrophic surface bacteria found in the Sargasso Sea, dominated by SAR11 clades I and II in BATS SUR waters [63,64]. As well, a higher proportion of viruses found in DCM samples were predicted to infect phototrophic *Prochlorococcus*, with *Prochlorococcus_B* the dominant *Prochlorococcus* ecotype and having greater than 6-fold abundance in DCM compared to SUR **(Fig 3A)**. This signal is consistent with *Prochlorococcus_B* being a low-light ecotype that prefers lower depths, like the DCM, at BATS [47,50].

**Figure 3.**
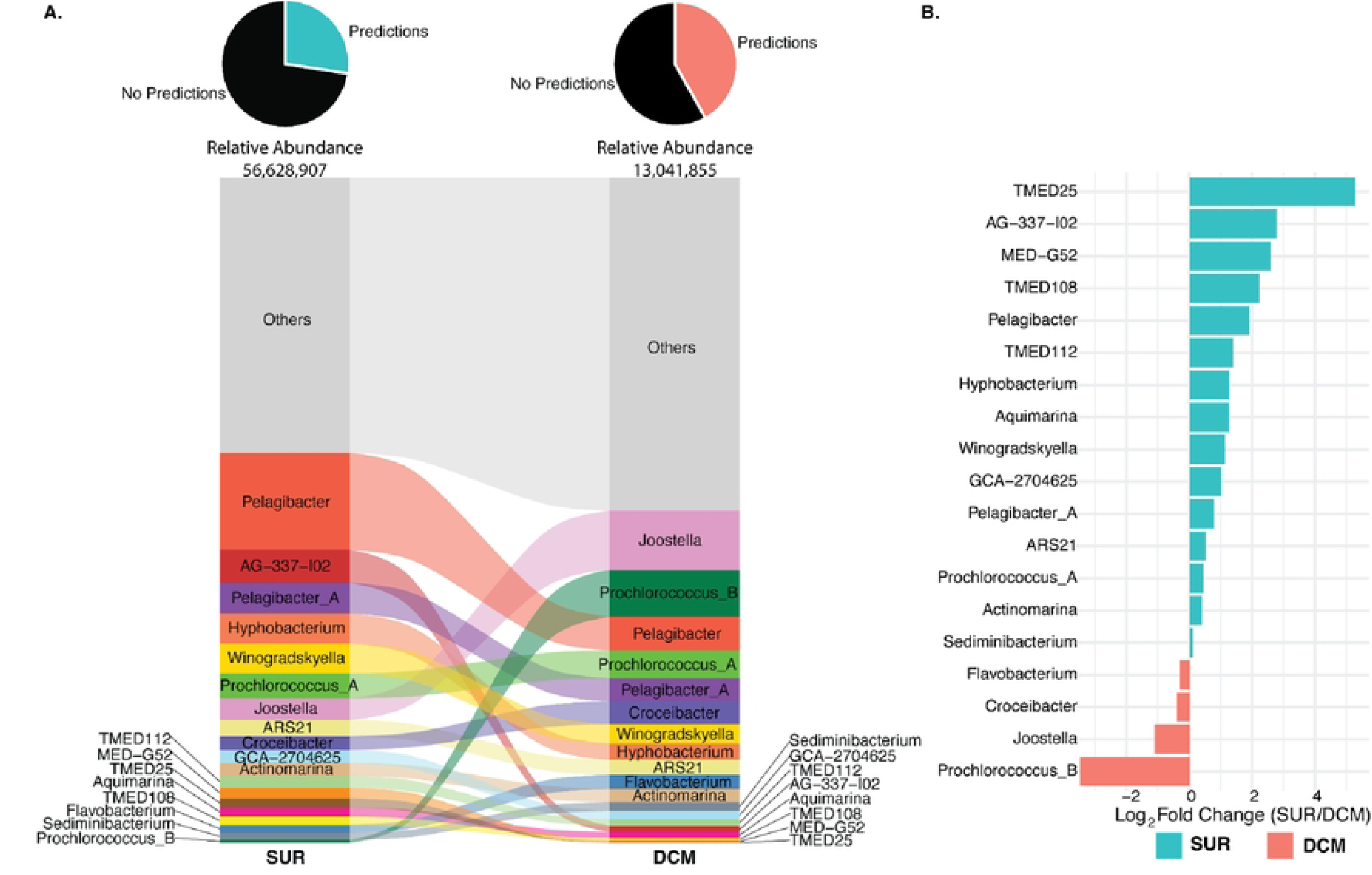
Viral population hosts across depths. **A.** Alluvial plot of predicted hosts (genus-level ranks) for viral populations present at each depth. Height of row is based on abundance of genera and genera that rnake up less than or equal to 1% of the total abundances are grouped as “Others”. Alluvial is ordered from 1nost abundant at top to least abundant at bottom. The pie chart on top of each column indicates the percent of viral populations with host predictions in each group with the total relative abundance of the category (SUR/DCM) below. **B.** Bar graph showing the Log_2_ fold change of host predictions across depths. Log_2_ fold change is based on abundances of genera that 1nake up greater than 1% of the total abundances in the dataset.

Looking further, we wondered whether viruses across these depths contained any metabolic genes of interest (e.g., auxiliary metabolic genes, or AMGs), and how these would fit into measured microbial metabolisms within the Sargasso Sea. [12–15,66]. We sought to uncover any AMGs in viruses unique to either depth environment. To ensure that genes were not misattributed as AMGs, we adopted a highly conservative approach for curating our AMG catalog, as used previously [52], which resulted in only ∼0.06% (399) of 684K annotated genes passing our conservative filtering. Beyond this conservative filtering, we did not conduct any manual curation or further validation.

Many of the putative AMGs belonged to pathways enriched in one depth over the other (**Fig 4)**. Among these, cobalamin biosynthesis was SUR-enriched and of particular interest given that *Prochlorococcus* and *Synechococcus* require cobalt for microbial growth [68,69] and comparative *Prochlorococcus* genomics has documented their genetic machinery for the synthesis and use of cobalt-bearing cofactors (cobalamins) [68]. Considering this, we hypothesized it was possible that abundant phototrophic viruses in the viral community harbor AMGs that assist in cobalamin biosynthesis. Assessing our data, however, we found a single gene in the cobalamin biosynthesis pathway (K09882, *cobS*) in each of 2 viral populations almost exclusively within SUR (**Table S6**). Host prediction suggested that one of these populations infect *SAR86A*, whereas the other had no predicted host. The *SAR86A* prediction is curious given that this lineage is not known to possess any genes involved in cobalamin biosynthesis.

**Figure 4.**
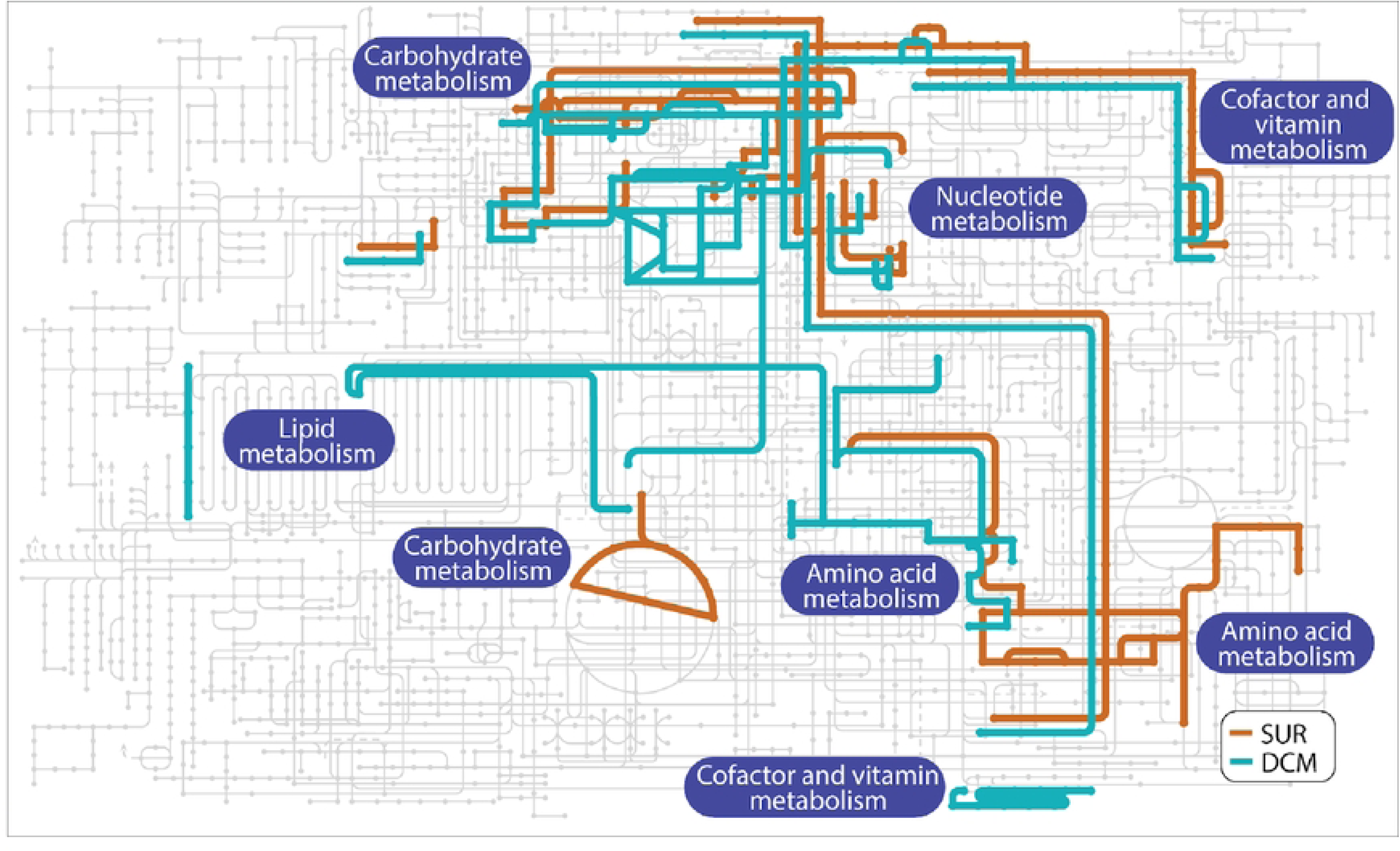
Metabolic path,vays found in viruses across depth gradients. KEGG 1nap was made on iPath (v3) under 111ap selection “Metabolic Pathways” using data fro1n Table S6 and shows pathways for AMGs present in viruses across depth gradients. Color of line is based on what depth the virus containing the AMG was found in, with gray lines indicating lack of AMGs relevant in that pathway for either group. This KEGG map is a qualitative analysis depicting which pathways identified AMGs belong to, with lines indicating pathways with relevant AMGs present.. Thickness of the line is the same throughout figure to highlight presence of pathways.

Taking instead, a host-focused perspective on AMG depth-related patterns, we observed the following. First, we explored virus populations predicted to infect *Pelagibacter* (formerly known as SAR11), i.e., pelagiphages [70], which revealed 4 AMGs among them including *phoH*, K06217 (2 viral populations) and 2OG-Fe(II) oxygenase, K07336 (2 viral populations). Of these, *phoH* is known to be present in both cyanophages [71] along with heterotrophic phages including pelagiphage [72]. Phylogenetic analysis of the *phoH* gene in phages has been shown to vary greatly based on the phage host while the role of *phoH* remains unclear as expression of the gene has been shown to vary within different phage-host pairs [73]. The 2OG-Fe(II) oxygenase AMG, however, is a much more well-known gene in pelagiphages [70]. Curiously, however, these pelagiphage AMGs were depth-specific with the viral populations possessing said genes only being present exclusively in one depth or the other. This can be seen with one of the viral populations predicted to target *Pelagibacter_A* that possessed a 2OG-Fe(II) oxygenase being found only in DCM, while all other pelagiphages with AMGs were found exclusively within SUR. Second, we explored AMG depth-related signals from *Prochlorococus*, which revealed a photosystem II gene (K02703, *psbA*) found in 7 viral populations exclusive to the DCM, where two populations were predicted to infect *Prochlorococcus_B* and the remaining five were predicted to infect *TMED112, TMED70, MGIIa-K1*, and 2 for unknown hosts. Within these host predictions both *TMED112* and *TMED70* are heterotrophic hosts while *MGIIa-K1* is a member of marine group II *Euryarchaea* [39]. *TMED112* is a member of the SAR86 cluster *Gammaproteobacteria* and along with *MGIIa-K1* are members of widely distributed and ecologically important clades of marine picoplankton [39,74,75], however, none of these genera have previously been shown to hold the photosystem II gene, psbA. Although both depths had virus populations with *Prochlorococcus_B* predicted as the host, this genus was a common (second most) predicted host in the DCM, 6x more abundant in DCM compared to SUR– all of which underscored its known low-light ecotype adaptations that would do well in DCM.

Finally, though a weaker signal, we highlight the finding that a SUR virus population contained a gene putatively involved in Coenzyme M biosynthesis (K05979, *comB,* 1 viral population), presumably relevant to methanogenic archaea. The coenzyme M biosynthesis signal might represent an understudied link to methanotrophic archaeal carbon metabolism, specifically anaerobic oxidation of methane to carbon dioxide [76,77]. Such methane cycling has previously been shown in oligotrophic North Atlantic Ocean gyres to be important with elevated methane from archaea as well as some bacteria that produce it as a byproduct [78]. Although the virus possessing this AMG had no host predicted, it is known that archaeal viruses outside extremophile conditions are incredibly challenging to predict [79]. We posit that the presence of a coenzyme M biosynthesis gene in a virus contig and its potential relevance to methane and carbon cycling at BATS, invites future targeted work once pelagic ocean archaeal viruses are more readily identifiable and/or increased genomic context becomes available for this gene.

### Diel periodicity analyses reveal population-level viral population dynamics

Though virus community-level alpha and beta-diversity metrics revealed no significant sub-daily temporal dynamics, we wondered whether specific populations had consistent diel cycles that might be detectable given the extensive sequencing available across this high-resolution time-course. We built upon previous work assessing diel periodicity in the ocean [80] to statistically detect diel patterns (via RAIN [81] and significance testing; see Methods) among our virus populations. This revealed 3,097 viral populations that were significantly diel rhythmic in SUR (10.68% of SUR viral populations). The lack of detected diel cycling viruses in DCM may be due to reduced sampling which may have limited statistical power to detect diel rhythmicity.

Focusing on SUR viral populations, we then compared diel versus non-diel SUR viral populations to look for differences at levels of host predictions, viral taxonomy, and gene function. For hosts, the abundances of viral populations for a given predicted host were summed and then assessed across the time course for diel versus non-diel viral populations via RAIN and significance testing as described above **(Fig 5)**. This revealed diel versus non-diel viral abundances for many predicted hosts with *Pelagibacter* being the most abundant targeted taxa across the two categories. However, slight differences were observed for the proportion of viruses targeting taxa like *Hyphobacterium* in the diel group or those targeting *Joostella* in the non-diel group. Such observations are difficult to interpret given that not much is known for either genus regarding their diel behavior in oceans. Similarly when looking at family-level viral taxonomy and their abundances, we observed very little change between diel and non-diel viruses, with slight differences in the enriched proportion of viruses belonging to Caudoviricetes (new family 33 and order 140) for diel and Caudoviricetes (new family 21 and order 134) in non-diel **(Fig S3)**.

**Figure 5.**
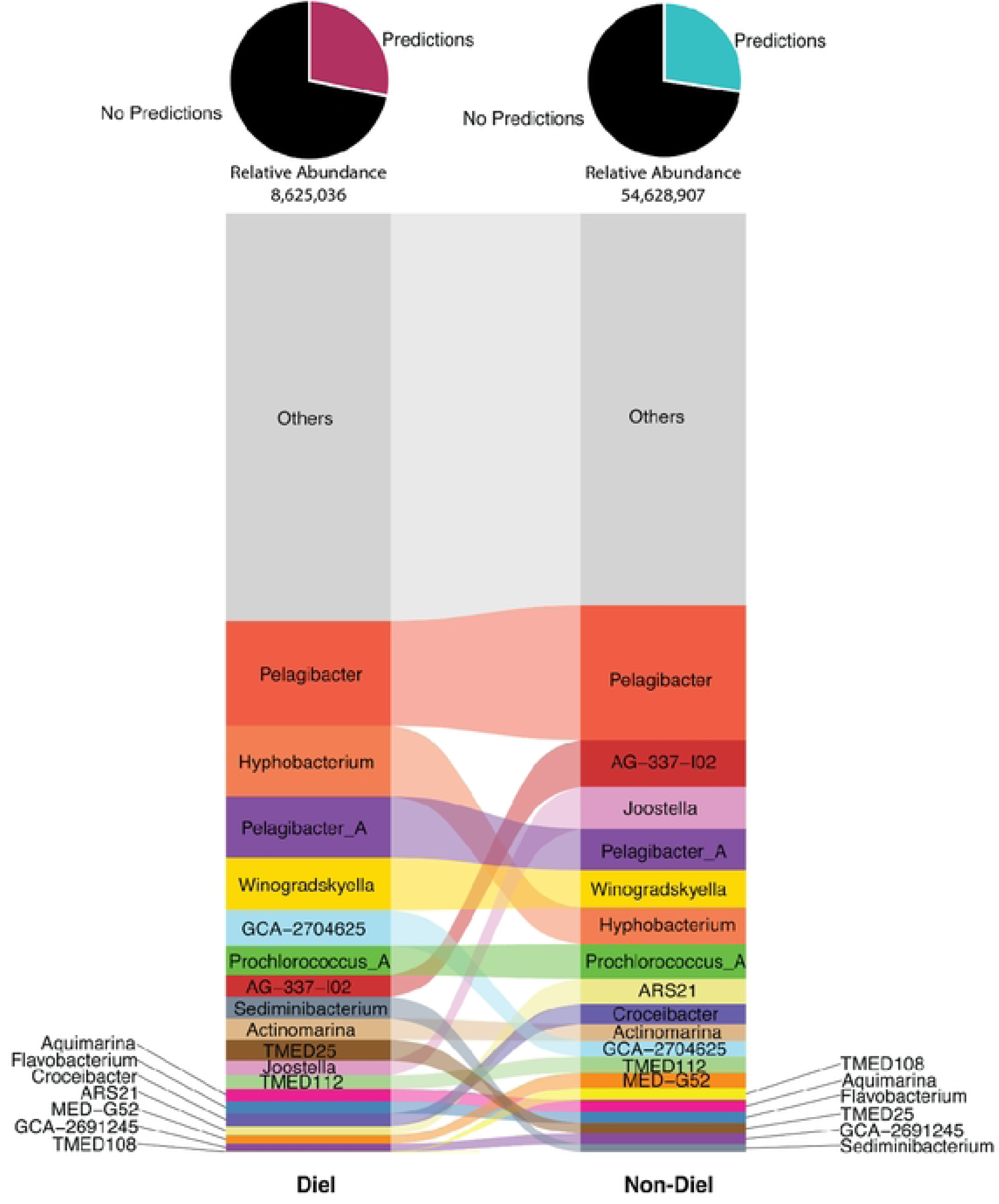
Predicted hosts for SUR viral populations,vith diet or non-diet rhythmic periodicities. Alluvial plot of predicted hosts (genus-level ranks) for diet and non-die! viral populations present. Height, order, and pie chart are as described in Figure 3.

Seeing largely similar host predictions and viral taxonomy for the diel and non-diel viral populations, we assessed their genes for functional gene content signatures. We hypothesized that viruses possessing diel patterns of abundance may possess AMGs that would enable these respective niches. Evaluating our conservatively filtered AMG catalog, we found 18 putative AMGs (belonging to 7 pathways) in diel viruses and 381 putative AMGs (belonging to 31 pathways) in non-diel viruses **(Table S9-10)**. Most of the genes identified by our conservative filtering methods for diel viruses were shared with non-diel viruses, while most of the non-diel virus genes were not. Among the AMGs shared between diel and non-diel populations, many are likely involved in virus infection to produce genomes and/or particles (e.g., nucleotide metabolism, amino acid metabolism). The only one that was exclusive to the diel viral populations was Coenzyme M biosynthesis (K05979, *comB,* 1 viral population), whereas the non-diel exclusive AMGs included 232 genes involved in 25 pathways (**Table S7-10)**. To summarize their biology, we interpret the non-diel-unique AMGs to represent pathways that contribute to constitutive rather than diel-cycling metabolisms. These constitutive metabolisms include genes involved in carbohydrate and redox metabolism (glucoronate, galactose, glyoxylate, ascorbate), nucleotide metabolism (purine, pyrimidine, degradation), amino acid and membrane metabolism (fatty acids, phospholipids), amino acid and polyamine metabolism (methionine, arginine/polyamines, creatine), cofactor and vitamin biosynthesis (THF, BH₄, B₁₂, biotin), sulfur cycling (sulfate assimilation/reduction) and secondary metabolites (aurachin). While less clear examples are those involved in cell wall/structural sugar biosynthesis (rhamnose, KDO) as dTDP-L-rhamnose biosynthesis which is involved in biosynthesis of terpenoids and polyketides is a pathway not typically found in viruses [82] (**Table S7-10)**.

### Unsupervised learning identifies day-versus night-peaking patterns in diel signals

We set out to evaluate the structure of diel rhythmic viral populations given observed variation in amplitude, shape of oscillation, and peak timing. To formally evaluate this, we built on recent efforts to identify viral archetypes in our dataset that may possess recurring temporal patterns, assign groups of viral populations to these archetypes, and assess within-and between-group differences in ecological relevance [80]. With this analytic pipeline [80], we utilized an unsupervised, self-organizing map (SOM) approach to determine whether smaller groups or ‘archetypes’ exist by evaluating the euclidean distances of their abundance across the time series (see Methods). The resultant SOM clustering approach for our 3,097 diel viral populations revealed 3 archetypes that we titled “archetype 1”, “archetype 2”, and “archetype 3” (**Fig 6A**). Adding to this archetype construction, we calculated the average peak time for individual viral populations as well (see Methods), revealing that most (94.09%) SUR diel viral populations peaked in abundance during the night with the most abundant time point being 04:00 hours, with much less common day peaking viral populations peaking at 12:00 hours.

**Figure 6.**
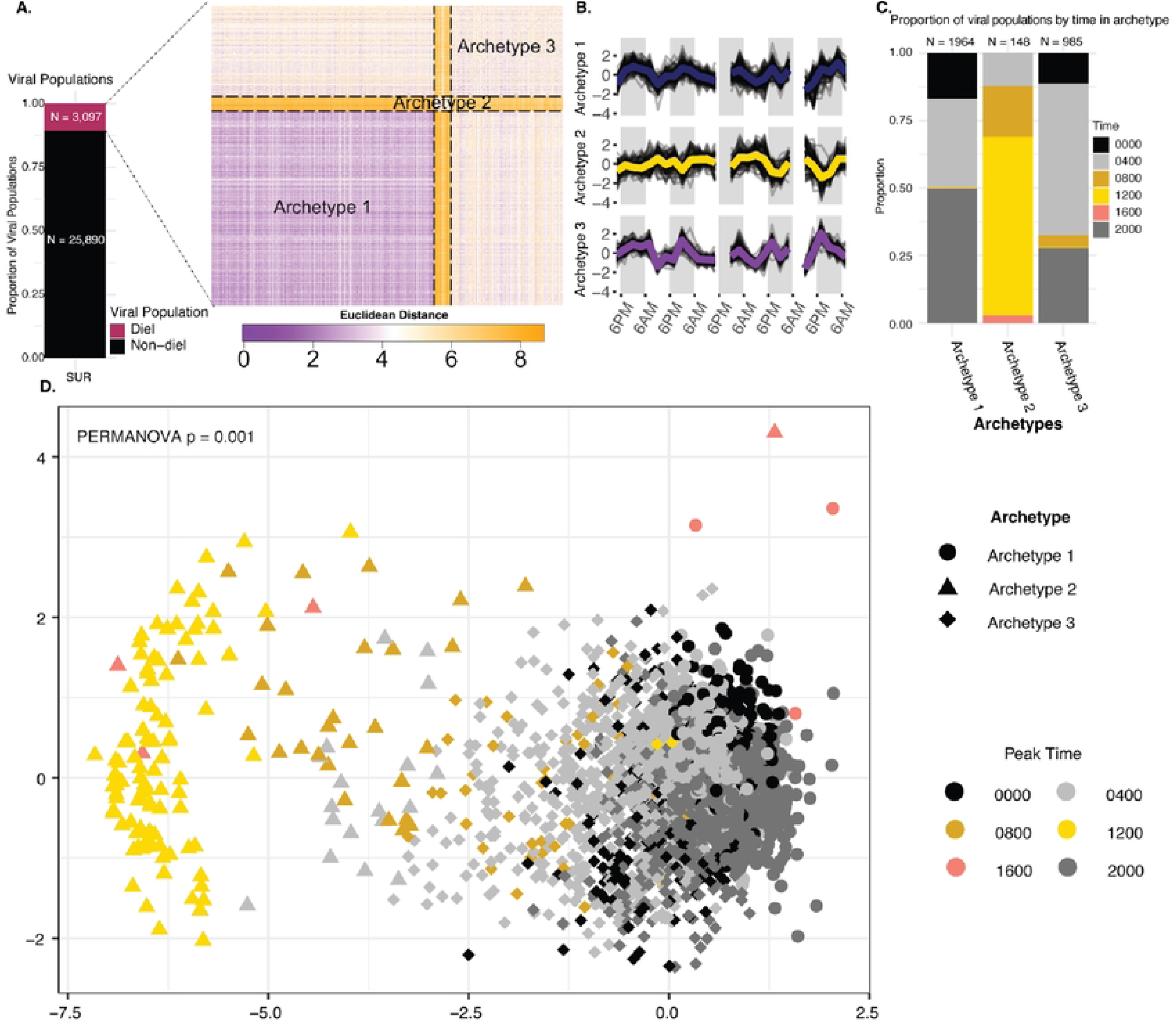
Categorizing diel patterns into three potential archetypes using unsupervised clustering. **A.** Organized pairwise distance 1natrix for all die! viral populations after clustering based on unsupervised, self-organizing maps (SOMs). Pixels in figure represent the Euclidean distance between the time series of two die! viral populations with purple indicating a sn1all distance (sin1ilar tune series) and orange indicating a higher distance (less similar time series). Dotted lines form boxes to separate the different cluster boundaries. **B.** Line graph for time series of viral populations within each cluster. Archetype tin1e series are a co1nbination of all ti111e series in their cluster as defined via the SOM algorithm (white line). A random sarnple of 145 tune series pertaining to each cluste are plotted as dark lines. C. Proportion for number of viral populations by tune in archetype. Colun111s represent archetype while color represents mean peak tune for viral populations across the tune series. **D.** NMDS of all die! viral populations colored by calculated peak rank 1neasuren1ent tune and given shapes by archetype.

To assess the archetypes for biological differences, we first evaluated their temporal patterns to connect those possible patterns to host predictions. These temporal patterning analyses revealed that archetypes 1 and 3 possessed night-peaking viral populations and dominated the dataset (95.22% of diel viral populations), whereas archetype 2 possessed primarily day-peaking viral populations and represented only a few percent of the total populations **(Fig 6B-C)**.

Ordination (NMDS with euclidean distances) revealed a clear separation between viral populations that coincided with time and archetype grouping such that one sees a gradual shift from virus population archetypes from peaking at 00:00 hours to peaking at 12:00 hours (**Fig 6D; Fig S4)**. Because these archetypes required no *a priori* assumptions of sinusoidal patterns nor of preferred phase, these observations represent naturally forming groups that presumably more accurately capture diel rhythmicity across the time course (see also [80]).

Evaluating these temporally classified archetypes further, we next sought to link these to changes in host predictions, which revealed dominant host prediction abundances varying between archetypes **(Fig 7)**. Archetype 2 consisted of viruses that predominantly targeted genera that made up ≤1% of the predictions in that archetype with only 4 different genera (*Winogradskyella, Pelagibacter, Joostella,* and *Flavobacterium*) having >1% of the host predictions. This is contrary to the other archetypes that possessed more host predictions that accounted for >1% of the viruses, perhaps because they were composed of so many more viral populations. Another notable observation was that archetype 1 held the largest proportion of phototroph targeting viruses, with viruses predicted to infect *Prochlorococcus_A* being the fifth most abundant host in the archetype **(Fig 7)**. Archetype 1 mainly consisted of viruses peaking in abundance during the night with the largest proportion peaking at 20:00 hours. Our observations are further supported by bacterial data sampled during the same expedition that show *Prochlorococcus* bacteria having larger abundances during nighttime samples [53]. The correlated abundance of viruses with their hosts seen here with *Prochlorococcus* coincides with other studies that used transcripts and have similarly seen phototrophic targeting viruses being co-expressed with their associated bacteria [40,41,83]. The night peaking viral activity of SAR11 and *Prochlorococcus* have also previously been observed as we did at BATS [40]. Our archetype comparisons, show that viruses that target heterotrophs are the main ones showing diel patterns and that among our archetype 2 viruses that mainly peak in abundance during the day, the biggest targets were the genera *Winogradskyella, Pelagibacter, Joostella,* and *Flavobacterium* **(Fig 7)**. As these genera are also found to be targeted in the other archetypes, it is possible that the difference here lies in a higher resolution of taxonomy for the bacteria being targeted.

**Figure 7.**
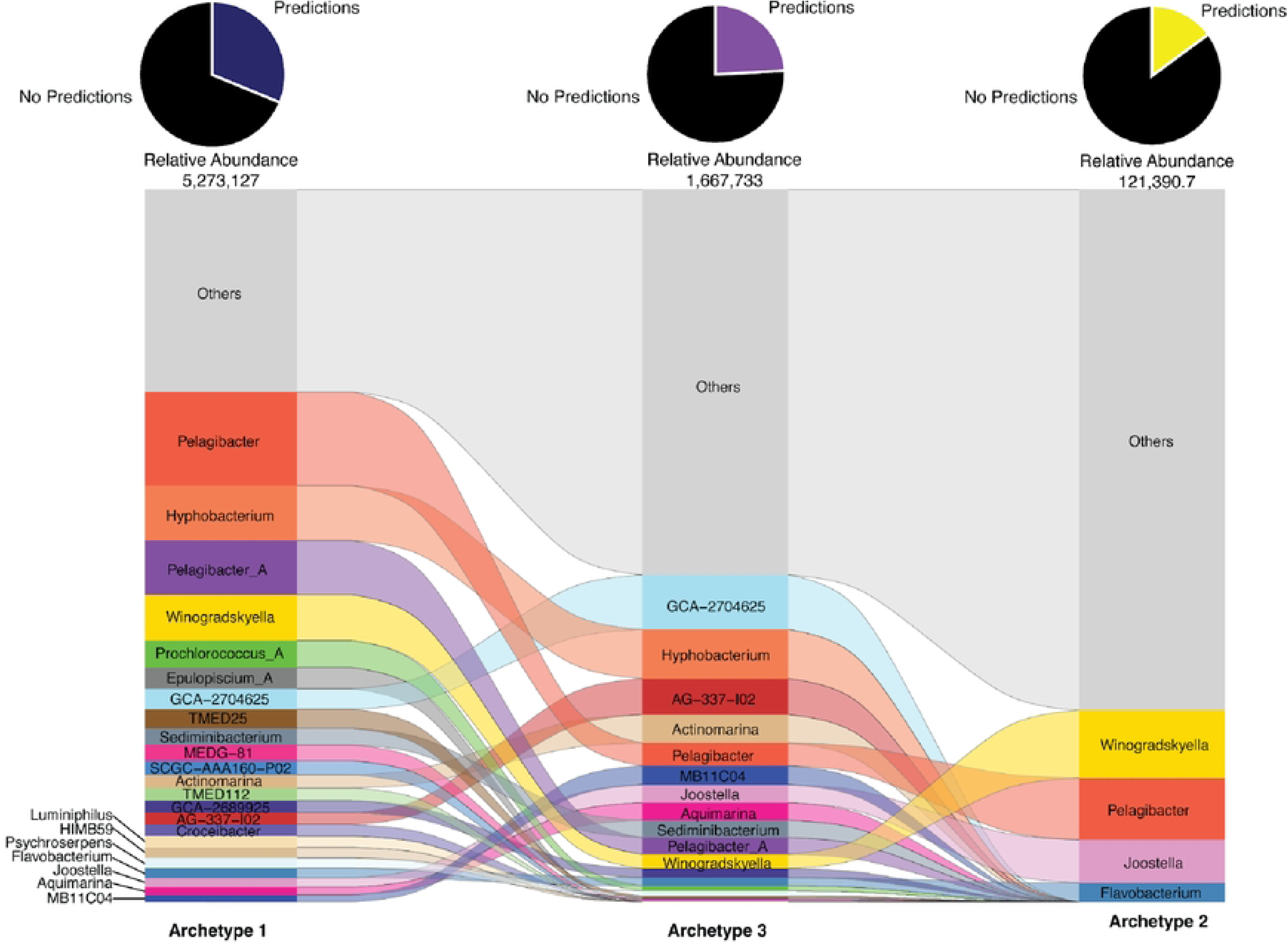
Comparison of diel archetypes host predictions. Alluvial plot of predicted hosts (genus-level ranks) for viral populations present within archetypes. Height, order, and pie chart are as described in Figure 3.

## Discussion

In this study, we leveraged a high-resolution sub-daily temporal dataset from SUR and DCM waters to investigate spatial and temporal viral dynamics at BATS, a model “future” ocean given climate predictions. With respect to depth, we observed distinct differences in viral alpha and beta diversity across the depth-gradient, which were likely shaped by physicochemical features inherent to each depth such as light, temperature, oxygen, and chlorophyll concentration [58–60].

Additionally, we expanded upon our prior work to show how host predictions for depth-structured virus populations changed at BATS [51]. Temporally, however, the picture was different – viral community-scale metrics remained stable across the 112 hour time course, whereas differences could be observed at the viral population-level. Exploration of these population-level differences revealed diel rhythmicity with machine learning-enabled methods [58] resolving such populations into one of three archetypes. Biologically, these archetypes differed in hosts predicted, viral taxonomy, and gene content.

By capturing viral abundances across depth and time, we uncover a complex and temporally structured viral community that mirrors host activity and environmental variation. The variation seen at the population-scale suggests that viral-host interactions are governed by more than abundance alone, likely shaped by biogeochemistry, host genome architecture, and ecological context [33]. Though not yet accomplished anywhere, future BATS efforts would benefit from many recent innovations that could layer in additional data, measurements, and interpretive frameworks. Long-read sequencing or single virus genomics could improve the capture of niche-defining genomic islands and microdiverse populations that are often missed by short-read assembled viromes [84,85]. Host predictions could be improved in a targeted manner through viral tag and grow experiments [86] or community-wide through DNA Hi-C proximity ligation sequencing, the latter assuming appropriate controls and analyses [87]. Measurements to assess activity, via metatranscriptomes, metaproteomes, metabolomes, and/or isotopic probing [88], could distinguish integrated past virus functioning emergent from viral particle sequencing from presumably real-time virus infection conditions. With better datasets, one could also expand the biological ‘players’ in the story, for example, by exploring the interplay between viruses and other mobile genetic elements to better inform the biological entities interrupting and/or carrying key metabolic reactions [89]. Additionally, modeling frameworks that predict viral roles in community metabolisms [90] and experiments that focus on phenotyping the ecological costs of resistance [17] could provide and test specific hypotheses about viral roles in biogeochemistry that will lead to better incorporation of viruses into predictive models. Again, such a diverse toolkit has not yet been applied anywhere, but does aspire to more holistically capture true biological integration across molecular and organismal scales that dictate ecosystem functioning.

On the whole, viruses impact marine ecosystems via lysis, gene transfer, and metabolic reprogramming during infection [91]. The BATS time series in the Sargasso Sea, with its multidecadal records of biogeochemical and physical data, provides opportunity to assess how viruses contribute to and are shaped by ocean warming, stratification, and shifting nutrient regimes [62], particularly as its gyre expands, warms, and acidifies [48]. Our work focuses on establishing baseline viral genomic datasets across two depths at BATS, where these observations provide glimpses into host-driven structuring of virus communities. Such observations show enrichment of viruses targeting *Prochlorococcus_B* and SAR11 clade bacteria in DCM and SUR respectively, along with genes predicted to be associated with said taxa such as *psbA* for *Prochlorococcus_B*. Along with this, we also document short-term temporal changes that manifest not in community-aggregated measures, but instead at the population-level where we revealed the diversity and potential functioning of viruses that differentially occupy day-or night-peaking niches. These short-term temporal changes not only illustrate that heterotrophic targeting viruses make the majority of our diel signal, but that 94.09% of these diel viruses peak in abundance during the night. Placing such short-term change into the context of longer-term changes already known for marine bacteria [63] will be critical to better incorporate viral ecology and roles into long-term ocean observing systems that seek to better predict future ocean functioning.

## Methods

### Sample Collection

Sea water samples were collected once every 4 hours in the surface, SUR or 12 hours in the deep chlorophyll maximum, DCM for 112 hours resulting in 39 samples (29 SUR, 10 DCM). A surface buoy with an underwater drogue, at 30 m depth, was deployed to allow us to follow the same parcel of water for the entire study. Water samples were collected using a CTD-rosette equipped with 24 x 12-l Niskin bottles. Depths for each cast were chosen based on in situ CTD oceanographic parameters. Sampling started at 16:00 local time (GMT-3) from SUR waters at ∼5 meters, always in the stratified SUR mixed layer, and 105-120m in the DCM mesopelagic layer. For each cast the CTD was deployed to at least 500 m to collect data on physical water columns, measuring the water temperature, pH, and salinity. Measurements were binned to the nearest 0.5 m of depth to standardize across casts, then a Nadaray-Watson kernel smoothing filter with bandwidth of 5 db was applied to each variable to remove noise and spikiness. For each sample, 10 L of seawater were filtered through a 0.2µm Millipore filter (CAT number: GPWP14250;LOG number ROBA92539) to minimize cells, and iron chloride flocculation [52] was performed to concentrate the viruses in the 0.2µm filtrate. The virus-concentrated iron hydroxide flocs were then filter-captured using a 1µm polycarbonate filter and stored damp on the filter at 4° C until resuspended in ascorbate-EDTA buffer and processed to DNA.

### Library preparation and sequencing

For each sample, half a filter, corresponding to ∼5L of seawater was resuspended in 7.5mL of ascorbate-EDTA resuspension buffer to capture the concentrated viral particles. All samples were then treated with DNase diluted 1:40 in DNase reaction buffer. DNase was then inactivated by adding 0.1 M EDTA then 0.1 M EGTA. Viruses were concentrated by spinning each sample through a 15mL 100kDa Amicon filter and resuspended in ascorbate-EDTA buffer (0.1 M EDTA, 0.2 M Mg, 0.2 M ascorbic acid, pH 6.0) in total of less than 1ml. DNA was then extracted using Wizard PCR Preps DNA Purification Resin and Mini-columns (Promega, Cat. #A7181 and A7211 respectively) [94]. Between 216 and 620ng of DNA (355ng on average between samples) was provided to the Joint Genome Institute for sequencing. Shotgun metagenomic sequencing at JGI was performed on an Illumina platform, resulting in an average of 144M reads per sample (∼9.8 x 10^7^ to 2.6 x 10^8^ sequencing reads per sample).

### Quality check, trimming, and assembly

Data processing and metagenomic analysis were conducted on the Ohio Supercomputer Center [95]. Reads went through a quality check using BbDuk (https://sourceforge.net/projects/bbmap/). Low-quality reads, adaptors, and Phix174 reads were removed (ktrim=r minlength=30 k=23 mink=11 hdist=1 hist2=1) and reads were trimmed (qtrim=rl maq=20 maxns=0 minlength=30 trimq=20). Four samples were resequenced due to low quality reads (T4D, T52S, T88S, T28D). Fastqc identified T28D as having a low read quality, so it was removed from the study.

Resequenced samples had an average sequencing depth of about 381M reads per sample, due to this, resequenced samples had their cleaned reads subsampled using bbmap 38.96 (https://sourceforge.net/projects/bbmap/) to an even depth of 62,381,078 in the SUR and 68,082,965 in the DCM which is the median depth value for clean reads in the rest of the samples in these regions. Samples were assembled using MegaHit [96] with default options.

### Viral Identification

Contigs from MegaHit were processed using Virsorter 2 SOP version 2.2.3 [54]. Briefly, all samples were first processed via Virsorter2 (–keep-original-seq –include-groups sdDNAphage, ssDNA –min-length 5000 min-score 0.5). Then CheckV v0.8.1 [97] was used along with Virsorter2 outputs to curate viruses following the parameters in the Virsorter 2 SOP [54]. After curation, the pipeline identified 228,013 viruses which were then clustered with CheckV clustering v0.8.1 [97]. 48,428 viral populations greater than 5kb and 14,634 greater than 10kb were clustered at 95% average nucleotide identity (ANI) and 80% coverage. A viral population is defined here as a group of viruses of the same species [46].

### Read mapping

CoverM v0.6.1 was used to map quality trimmed reads to viral populations (https://github.com/wwood/CoverM; –output-format dense, min-read-percent-identity 0.95, min-read-alignment-percent 0.75, min-covered-fraction 0.75,-m trimmed_mean). Coverage values were normalized by dividing them with read depth from the quality trimmed reads per sample and multiplying by 1,000,000,000.

### Diversity calculations and statistical analyses

Diversity estimates were based on the normalized relative abundance tables generated via read recruitment. The alpha (Inverse Simpson) and beta (Bray-Curtis dissimilarity) diversity statistics were calculated using vegan on R (v 2.6.4) [98]. Read were log2 transformed before calculating Bray-Curtis (function vegdist, method = ‘bray’). Affinity propagation using APCluster on R was used to cluster samples into groups with similar viral relative abundance and community structure [99]. Communities were then divided into groups based on similarity for analysis and on depth and time of day for separate alpha/beta diversity analysis. Principal coordinate analysis (function cmdscale) and nonmetric multidimensional scaling (function metaMDS, permutations = 999) were ordination methods used on the Bray-Curtis dissimilarity matrices [100–102]. The statistical significance between viral communities was validated by comparing the within community and between community distances with MRPP and ANOSIM [103,104].

### Temporal analysis with the detection of rhythmic behavior in time series

For all datasets from BATS, diel periodicity was determined using the rank-based Jonckheere-Terpstra umbrella test implemented in the R RAIN package [81]. Normalized read abundance tables of the viral populations were observed for oscillating behavior across the 112-hour time series. Samples were first detrended (linear regression with respect to time was subtracted from time series) to increase power of rhythmicity detection using the detrend function in the R pracma package v.2.4.4 [81]. RAIN was run with the options method = ‘deltat = 4, period = 24’ for the surface samples to establish four-hour sampling periods and a 24-hour time period to observe for oscillation. Viral populations within the SUR samples had the options method = ‘measure.sequence = c(1,1,1,1,1,1,1,1,1,1,1,1,1,0,1,1,1,1,1,1,1,1,0,1,1,1,1,1,1)’ to account for sample “T52_S” and “T88_S”, that were excluded from analysis. Samples from the deep chlorophyll maximum (DCM) were run with the options method = ‘deltat = 12, period = 24’ to establish 12-hour sampling periods and a 24-hour time period to observe for oscillation. Viral populations within the DCM samples had the options method = ‘measure.sequence = c(0,1,0,1,1,1,1,1,1,1)’ to account for samples “T4_D” and “T28_D” that were excluded from analysis. After RAIN implementation, the Benjamini-Hochberg false discovery rate (FDR) control procedure was implemented to assess significance at the P = 0.05 level for each data type.

### Calculation of peak rank time

To estimate the mean peak time for viral populations, a rank-based heuristic was calculated. For a given viral population, the abundance at each time point was ranked. The ranks from all measurements were then averaged and the peak mean rank time was defined as the time with the highest average, where ties were summarized as the center between tied times [80].

### Clustering Analysis

Detrended diel time series were scaled to make data dimensionless to reduce impact of magnitude on euclidean distance matrices. Euclidean distance matrices had a Hopkins statistic calculated to determine the meaningfulness of clustering where a value of *h = 0.71* was found, indicating data structure cannot be explained by random distribution of distances. To determine the best clustering method we employed hierarchical clustering (implemented by the hclust function in the R stats package v.4.1.1), medoid clustering [105], and training of SOMs [106]. These clustering methods were then evaluated for best fit using the Calinski-Harabasz metric and average silhouette distance. SOM was selected as the best fit clustering method on the basis of identifying the ‘elbow’ in decreasing average silhouette width to initially select three as the operational number of clusters. Fits were assessed in more detail for two and four clusters as well.

### Taxonomic Classification

Prodigal v2.6.3 [107] was used to predict proteins from the viral populations greater than 10kb. From the 15,197 viral populations greater than 10kb, prodigal predicted 360,993 proteins. vConTACT 2 version 0.11.1 [108], was used to classify viral genomic sequence data. This specific tool is designed to cluster and provide a taxonomic context of metagenomic sequencing data. VConTACT2 results were then read into cytoscape v3.10.1 for visualization of the network following the protocol published on protocols.io [108].

### Functional annotation and auxiliary metabolic genes (AMGs)

AMGs were annotated using DRAMv v1.4.0 (min-contig-size = 10000, --skip_trnascan). AMGs with a score of 1 to 3 were selected as putative AMGs [109]. AMGs were then curated for a conservative catalog [67] then filtered for those containing pathway information from KEGG. The conservative AMG catalog was mapped to metabolic pathways using Anvi’o v8 [110].

Metabolic pathways were visualized in KEGG Metabolic Pathways (map 01100) with iPath 3.0 [111] based on the presence of AMG-carrying viruses across depth layers and their diel signals.

Pathways with default stepwise completeness less than 0.75 were considered incomplete.

### Identifying hosts using iPHoP with the standard database

IPHoP v1.3.2 [62] was used to predict hosts from viral populations >5kb. Hosts were first identified using the standard database using the default commands.

### Identifying hosts using iPHoP with a custom database

A custom database was also created using 89 metagenome-assembled genomes (MAGs) from a previous BATS study done in July of 2017 [51]. Following the protocol on iPHoP’s github page (https://bitbucket.org/srouxjgi/iphop/src/main/), GTDB [112] was used to assign taxonomy to the MAGs, which were then added to the standard iPHoP database to create a custom database.

GTDB was run separately identifying bacteria with the outgroup *p_Patescibacteria* and archaea with the outgroup *p_Altiarchaeota*. Identified MAGs were then added to the standard iPHoP database using the add_to_db command. Identifying hosts from the custom database was done using the predict command.

### Sample collection from which MAGs were identified

The custom database uses MAGs identified from Warwick-Dugdale, 2017 dataset [51]. These MAGS were derived from 12 samples at the Bermuda Atlantic Time series (BATS) station (31°40’N, 151 64°10’W) during dusk (∼19:00 local time) and dawn (∼06:00 local time), from the depths of 80m and 200m, over a period of four consecutive days from the 8^th^-11^th^ of July 2017.

## Acknowledgements

The authors would like to thank the captain and crew of the RV *Atlantic Explorer* for conducting cruise AE1926 along with the marine technicians who helped in the collection of seawater samples. The authors would also like to thank Garrett Smith, Rokaiya Shatadru, James Riddell V, and James Tan for their suggestions to the research and manuscript, along with the OSU Microbiome Informatics course Microbiology 8161 for their support in the early stages of the analyses.

## Funding

We also acknowledge the commitment and support provided by the CMBP grant NIH T32 GM141955. This study was partly supported by iVirus2 (award number 2149505), iVirus2 (award number NSF ABI#2149505), VirSoil2 (award number DOE#DE-SC0023307), and InVirT (award number NSF OCE-1829640 and OCE-1829641). JSW acknowledge support of NSF grant OCE-1829636. J.S.W. is an investigator at the University of Maryland-Institute for Health Computing, which is supported by funding from Montgomery County, Maryland and The University of Maryland Strategic Partnership: MPowering the State, a formal collaboration between the University of Maryland, College Park and the University of Maryland, Baltimore.

## Data Availability

Raw viromic sequencing data is available through the JGI under project ID #505733. All supplemental tables and large supplemental data can be found at https://doi.org/10.5281/zenodo.17260609. Statistics and figures code base are available on a GitHub repository https://github.com/carrillo1998/BATS, please contact MBS for any further data or code requests.

## Notes

### Competing Interest Statement

The authors have declared no competing interest.

